# Graft protection by myeloid progenitor cells using lethal or non-lethal preconditioning

**DOI:** 10.1101/2025.11.25.690526

**Authors:** Yongwu Li, Lei Sun, Erica Martinez, Anna Sedello, Timothy Fong, Jos Domen

## Abstract

Myeloid progenitor cell (MPC) therapy can protect against infection and can be used as an irradiation countermeasure, but can also prevent rejection of matched grafts. Here we report on the matching requirements between MPCs and skin grafts, and on the development of a non-lethal preconditioning model that supports MPC-induced protection of skin grafts. Mouse MPCs (mMPC) were obtained from HSC following 10-day *ex vivo* expansion. Skin grafts were tested in host mice that either received a lethal dose of irradiation and reconstitution with allogeneic mMPC and third party HSCs or were given injections with depleting antibodies and chemotherapeutics followed by transplantation of mMPC without HSC. mMPC-matched grafts were protected and no significant difference was observed between mice receiving irradiation and fully matched grafts (major and minor transplantation antigens), partially matched grafts (major only), and fully matched grafts in mice that received sublethal irradiation. Haploidentical grafts (half of the MHC alleles not matched at all) are not protected. We conclude that closely matched mMPCs are sufficient to prevent rejection. We also show that mMPCs are effective in a trachea transplant model. Graft protection does not require either lethal preconditioning or an accompanying HSC transplantation, and affects both T- and B-cell responses.

## Introduction

In organ transplantation acute rejection is well controlled, but long-term organ survival remains more problematic. Chronic rejection is a complicated process that is not fully controlled [1, 2]. Long-term use of immunosuppressants has many associated morbidities, including neural and renal toxicity and increased risk for cancer and infections [3–6]. Teaching the otherwise functional host immune system to not reject the transplanted organ, to be tolerant of it, would abrogate the need for constant immunosuppression. However, the complexity of the immune system has made this difficult to accomplish.

Cell-based therapeutic approaches aimed at modulating the recipient’s immune response to achieve the above mentioned goal are of interest in increasing long-term organ survival [7]. These include regulatory T cells, dendritic cells (DCs), regulatory macrophages, and myeloid derived suppressor cells. Many cell-based approaches are being tested pre-clinically and clinically, e.g. in the context of the ONE study [8–12]. Our efforts focus on the use of myeloid progenitor cells (MPCs). Unlike more mature cells, MPCs can differentiate into all (immunomodulatory) myeloid cell types, and they can expand extensively after transfer [13, 14].

MPCs have been developed into a clinical product based on their ability to reduce infectious complications under neutropenic conditions [15, 16]. Preclinical studies have shown that they can be used as a radiation countermeasure using either our [17] or alternative expansion conditions [18]. Of particular significance for clinical application: MPCs can be expanded *ex vivo*, can be cryopreserved before use, and are functional without the need for haplotype matching in the context of neutropenia [14]. A cryopreserved, allogeneic human MPC product (romyelocel-L) is being developed to prevent bacterial and fungal infections in patients with prolonged neutropenia and has been evaluated in three clinical trials to date (NCT00891137, NCT01297543 and NCT02282215). We have established that mouse MPCs (mMPC) administered at the time of HCT/skin graft placement will prevent rejection of matched skin graft transplants. Mouse studies with Rag2-/-Il2rg-/-mice as mMPC-donors have proven that skin graft acceptance does not require graft-matched lymphoid cells [13, 19]. Similar results were recently reported in independent studies [18]. In this paper we define the presence of allogeneic skin grafts for more than 5 months as tolerance, while the ability of these animals to reject non-matched skin grafts as evidence for a functioning, if not necessarily normal, immune system after preconditioning.

In order to further develop this approach, we are defining parameters important for clinical application of graft protection by MPCs. Specifically, we present data on the degree of transplantation antigen matching required between mouse MPCs derived by *ex vivo* expansion (mMPC) and skin grafts. We also demonstrate that protection by mMPC extends to a second transplant model: Heterotopic trachea transplantation. In addition we have developed a non-lethal preconditioning model that allows for mMPC administration 1 or more weeks after skin graft placement. We will discuss these results, and the effect mMPC administration has on T and B cell responses in these mice.

## Methods

### Mice

Female mice were purchased from Jackson Labs (Fig 2E). FVB mice, from which HSCs were isolated for expansion into mMPCs, were purchased from Charles River. The animals were maintained and used at University of Missouri Kansas City LARC or Cellerant Therapeutics under IACUC approved protocols. Donor mice were 5-8 weeks (HSC) or 8-12 weeks of age (mMPC); recipients were 7-12 weeks old.

**Figure 1.**
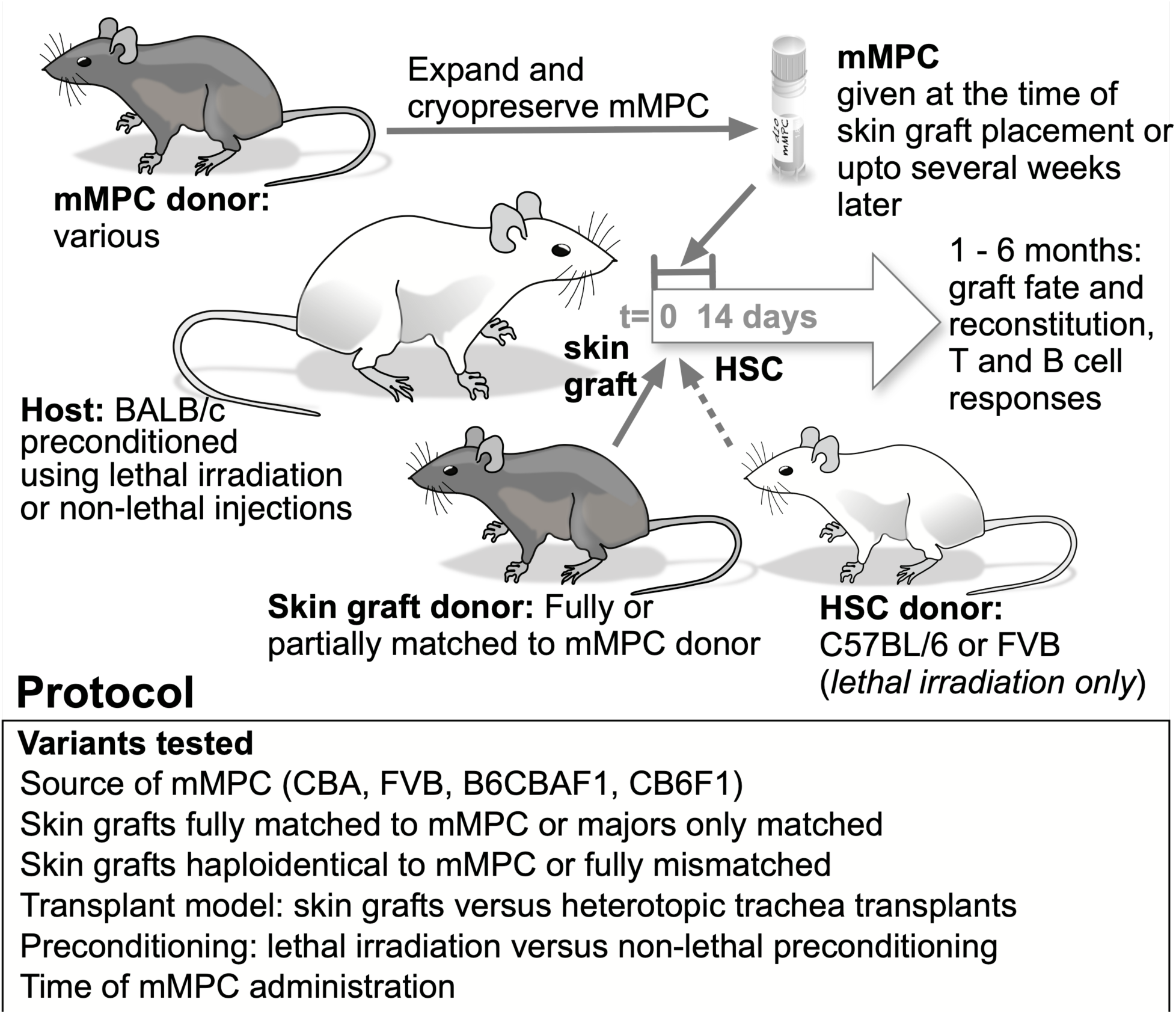
(A) Overview of the experimental design. Lethally irradiated mice were reconstituted with 4×10^3^ to 6×10^3^ HSC and 5×10^5^ day 10 expanded mMPCs. Skin grafts were placed at the time of irradiation and HCT. Non-lethally preconditioned mice were reconstituted with 3 to 6×10^6^ day 10 mMPC. Skin grafts were placed 1 or 2 weeks before mMPC administration (see Fig 7A for more detail). Skin graft fate and hematopoietic reconstitution was followed for 6 months, trachea grafts were analyzed after 1 month. In some animals T and B cell responses against mMPC matched and mismatched targets were determined repeatedly during the first 6 months.

**Figure 2.**
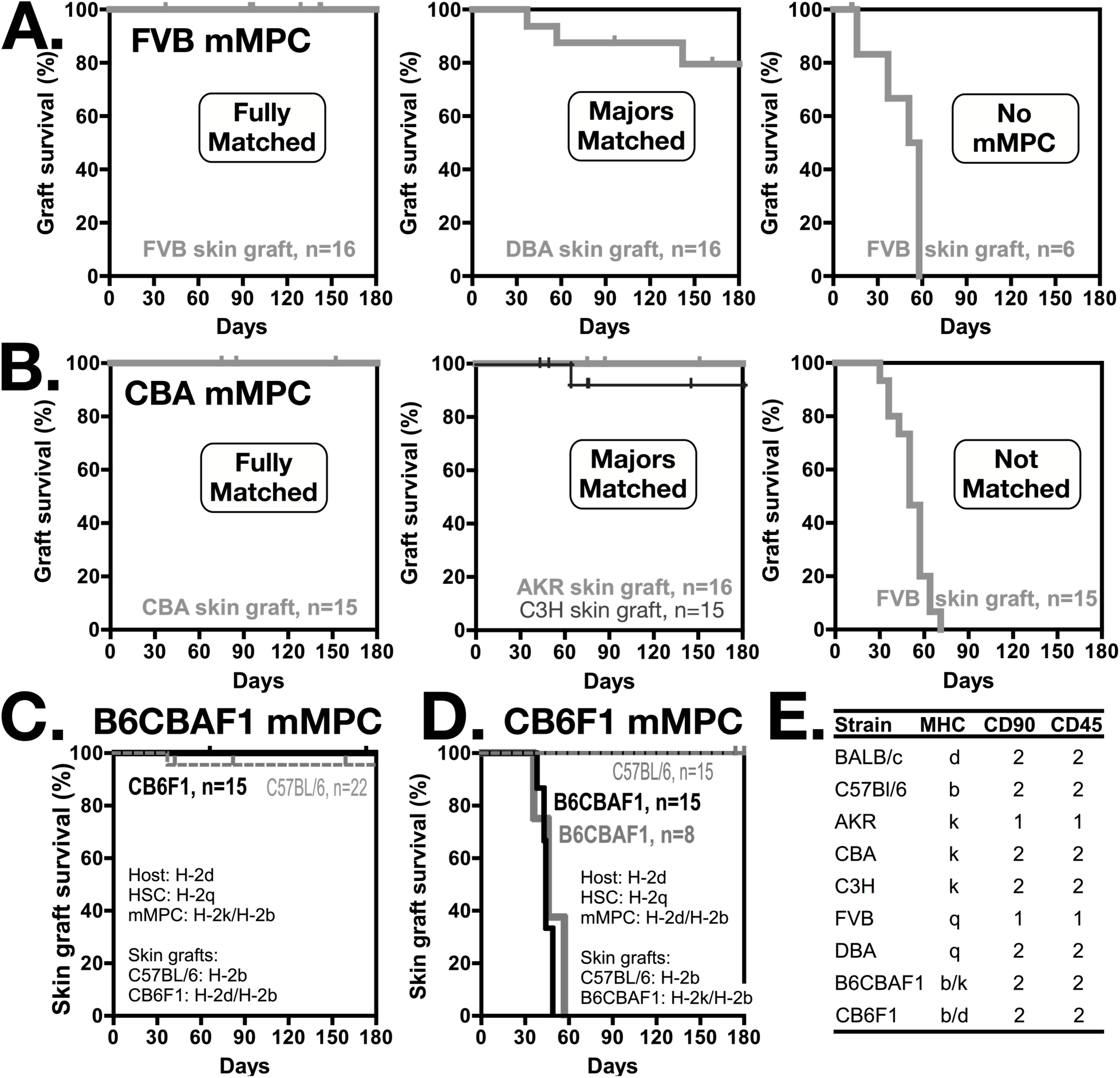
Skin graft survival using fully and partially matched mMPC. The plots show 6 months skin graft survival data for BALB/c (H-2d) mice reconstituted with 5×10^5^ day 10 mMPCs from (A) FVB (H-2q) or (B) CBA (H-2k) mice. Skin graft matching to the mMPCs was performed as indicated in the plots. The mice also received 6×10^3^ C57Bl/6 (H-2b) HSCs. Skin grafts were placed at the time of HCT and irradiation. (C) A heterozygous skin graft from a CB6F1 (H-2d/H-2b) will be protected if one set of MHC alleles (H-2) is present in the B6CBAF1 mMPCs (H-2b) and one in the host cells (H-2d) (black curve). Grafts from C57BL/6 (H-2b) mice were protected since the MHC antigen was also matched to the B6CBAF1 mMPCs. (D) Heterozygous skin grafts from B6CBAF1 mice (H-2b/H-2k) that contain one set of MHC alleles (H-2k) not matched to the either the CB6F1 (H-2d/H-2b) mMPCs, HSCs (FVB, H-2q) or host (BALB/c, H-2d) are rejected (black curve) with similar kinetics as non-matched grafts. Increasing the mMPC dose to 3 million cells (grey line) does not improve outcome. Survival of the C57Bl/6 skin grafts (thin dashed grey line) is a control to show that the CB6F1 mMPCs are functional. BALB/c mice received 5×10^5^ day 10 mMPCs and 4×10^3^ FVB HSCs at the day of skin transplant (C and D).

### Preconditioning

For lethal preconditioning the BALB/c recipients received 8.5 Gy at 1.2 Gy/min in two doses three hours apart (Radsource, RS 2000 x-ray machine, UMKC). For non-lethal preconditioning the mice were injected as shown in Fig 7A with 20µl anti-thymocyte serum per mouse (ATS) (Accurate Chemicals, NY) followed by repeated injections with 2.5mg/kg anti CD3 (clone 145-2C11, BioXCell, NH) and single injections with busulfan (10mg/kg) and rapamycin (24mg/kg) (both Sigma-Aldrich, MO). Preconditioned mice received antibiotic water (10^6^ U/L polymyxin B sulfate and 1.1 g/L neomycin sulfate) for four weeks (Sigma-Aldrich, MO).

### Derivation of mMPC

mMPC were derived from sorted HSC as described [17] using a 10 day expansion protocol in X-VIVO15 medium supplemented with 50 ng/mL recombinant mouse SCF, 5 ng/mL recombinant mouse TPO (R&D Systems, Minneapolis, MN), 30 ng/mL recombinant mouse Flt3L (Invitrogen/Thermo Fisher Scientific, Waltham, MA), 100 µg/mL Primocin (InvivoGen, San Diego, CA), and 1x Glutamax (Invitrogen/Thermo Fisher Scientific). Mouse MPCs were cryopreserved in formulation buffer comprising of CSB (BioLife Solutions, Bothell, WA) supplemented with 8.5% DMSO and 5% human serum albumin at approx. 2×10^7^ cells/mL.

### Transplantation of mMPC

Frozen cells were thawed in a 37°C water bath. Post-thaw viability and cell counts were determined by Trypan Blue (Gibco/Thermo Fisher Scientific, Waltham, MA) exclusion. mMPCs were resuspended at the desired concentration and, in lethally irradiated mice, were combined with freshly sorted HSC for transplantation into anesthetized mice by retro-orbital intravenous injection. For mitotic inactivation the mMPC were incubated for 30min with 100µg/ml of mitomycin-C (Sigma-Aldrich, St Louis, MO) prior to injection.

### Flow cytometry

HSCs were prepared as described [17, 19] using cells stained for CD117^+^Lin^neg/lo^Sca1^+^ (KLS) or CD117^+^CD90.1^lo^Lin^neg/lo^Sca1^+^ (KTLS). Cells were sorted using a FACSAria (BD, San Diego, CA) operated by the Flow Facility Core of Kansas University Medical Center.

For analysis (bone marrow, spleen, thymus or blood) erythrocytes were sedimented using 2% dextran (Pharmacia) in PBS (blood only) and remaining erythrocytes were lysed with 0.15M ammonium chloride/0.01M potassium bicarbonate. The cells were blocked with rat IgG (Sigma), divided into aliquots, and stained with antibodies as indicated, using MHC class I and CD45-allelic markers to distinguish origin. Antibodies used to distinguish cell origins (Table 1) included H-2b (AF6-88.5), H-2k (36-7-5) (both Biolegend, San Diego, CA), H-2q (KH114), H-2d (34-2-12) (both BD) and CD45.1 (A20) (eBioscience). Lineage markers were CD45R/B220, CD3, CD4, CD8, CD11b, and Gr-1 as indicated above. The cells were analyzed using an Attune flow cytometer. B cells are defined as CD45R^+^ (B220), but negative for CD4, CD8, Gr-1 and CD11b. To determine anti-H-2 antibody levels 1 million target splenocytes (as indicated) were incubated with serum (1:15 dilution), followed by incubation with rat-anti-mouse-kappa-FITC and anti-CD3-APC. CBC counts were obtained using a ScilvetABC veterinary blood cell counter using mouse presets.

### Mixed lymphocyte reactions (MLR)

Blood was collected and WBC isolated as described above. 2.4×10^4^ Celltrace CFSE (Invitrogen) labeled WBC were mixed with 3.6×10^4^ mitomycin C treated stimulator cells (C57Bl/6 or FVB) in triplicate in a 12µl volume per well in Terasaki plates. The cells were incubated for 3 days. Controls included no stimulator cells and stimulation with 2µl/ml CD3/CD28 beads + 30 u/ml IL-2. Cells were stained (H-2d-PE, H-2b-PE-Cy7, CD45.1-APCCy7 and CD3-APC, see above for clones) and the percentage of CFSE low staining cells for the H-2d+ CD3+ cells was determined.

### Transplants

Skin transplants mice were performed as described [13, 19]. Briefly, mice were anesthetized, tail skin was removed from donor mice and secured on the belly of host mice using interrupted stitches. Grafts were protected the first two weeks using band aids. Grafts were documented using regular digital photography. Skin graft sizes were calculated from these photos using an embedded scale.

For heterotopic trachea transplants tracheas were removed from donors and inserted lengthwise in a subcutaneous pocket in between or just below the scapulae. The skin was closed with 6-0 Maxon or Biosyn sutures. Three tracheas were placed on each host. The transplanted tracheas were removed for histological analysis after one month.

### Statistical Analysis

Data were analyzed using Graphpad Instat 3.0 and Graphpad Prism 4.0 (Graphpad Software Inc, San Diego, CA). Students T-test and Kaplan Meier with a Wilcoxon logrank test for groups was used. A p<0.05 was considered significant. Results are shown as means ± standard deviations.

## Results

### Transplantation of Major MHC matched mMPC

mMPC can protect fully matched, but not completely mismatched skin grafts [13, 19]. Here we tested skin grafts that were matched for the major MHC loci, but not the minor transplantation antigens. mMPC were generated and cryopreserved using the 10 day expansion protocol. BALB/c mice received lethal preconditioning and a transplant of 4×10^3^ to 6×10^3^ FACS purified HSCs, fully mismatched to the host, and 5×10^5^ mMPC fully mismatched to the host and HSCs.

Zero out of 31 skin grafts placed were rejected from animals that received mMPCs that were fully transplantation antigen-matched to the skin grafts, as illustrated in Figures 2A-B and Figure 3. Some graft loss (4 out of 47 grafts were rejected) was observed in the six-month observation period in the animals that received grafts matched to the major but not minor mMPC transplantation antigens. The difference is not statistically significant, p=0.1 (logrank), not even when the fully matched grafts from the experiments shown in Fig 2c and d are added, bringing the total 6 month graft survival for fully matched grafts to 82/83 mice, p=0.07 (logrank). All grafts were rejected from animals that either received preconditioning and skin grafts not matched to the host, HSC or mMPC donor or did not receive mMPCs.

**Figure 3.**
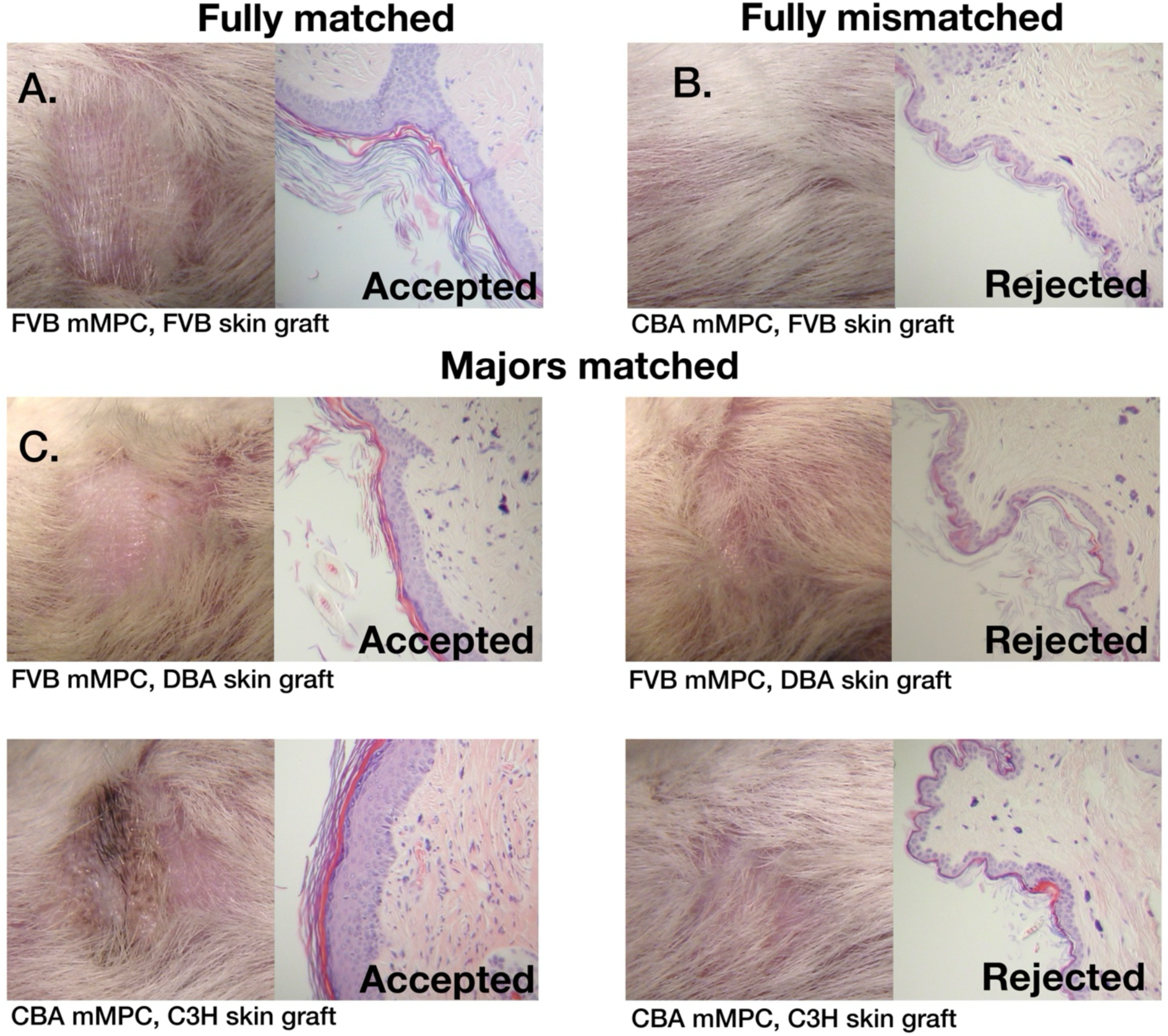
Histological analysis of skin grafts. Examples of (A) an accepted skin graft (FVB skin grafts in BALB/c hosts reconstituted with C57BL/6 HSCs and FVB mMPCs) harvested 214 days after placement and (B) a rejected skin graft (FVB skin graft on BALB/c hosts reconstituted with C57BL/6 HSC and CBA mMPCs). Analysis was performed 124 days after placement of the graft (which was rejected in 71 days). (C) Illustrations of skin grafts matched only to the major transplantation antigens of the mMPC. The middle row shows DBA skin grafts placed on BALB/c hosts receiving C57BL/6 HSC and FVB mMPC, 215 days after graft placement. The bottom row shows C3H skin grafts placed on BALB/c mice receiving C57BL/6 HSC and CBA mMPC, 234 days after placement. Accepted grafts (left panels) show the typical thick epithelium layer of the transplanted tail skin while the two rejected grafts (right panels) have reverted to the thin epithelium of normal trunk skin.

### Skin graft acceptance for haploidentical grafts

We also tested the ability of mMPCs to protect haploidentical skin grafts using F1 mice as skin and mMPC donors. In control experiments the host mice were BALB/c (H-2d) while the HSC donors were FVB (H-2q). B6CBAF1 mMPCs (H-2b x H-2k) protected both C57BL/6 skin grafts (H-2b) as well as CB6F1 skin grafts (H-2d x H-2b) (Figure 2C). The H-2b haplotype is present in the mMPC while the H-2d haplotype is present in the host mice. When testing haploidentical mMPC, CB6F1 mMPCs (H-2d x H-2b) fully protected control C57BL/6 (H-2b) skin grafts, confirming that the CB6F1 mMPC are functional. However, CB6F1 mMPCs (H-2d x H-2b), used as 5×10^5^ cells per mouse (n=15) or 3×10^6^ cells per mouse, (n=8) did not protect B6CBAF1 (H-2b x H-2k) skin grafts with 0/23 grafts surviving past 60 days. The k haplotype in the B6CBAF1 skin grafts is not matched to either host, mMPC or HSC (Figure 2D). This demonstrates limits to the ability of mMPCs to protect grafts.

### Reconstitution analysis in mice given HSC and mMPC

As reported before, the majority of circulating cells are HSC-derived, with minor contributions from mMPC- and host-derived cells [13, 19]. mMPC contribution is strongest in the first month (Figure 4A,B). CB6F1 mMPC show similar kinetics, but higher reconstitution levels (Figure 4B).

**Figure 4.**
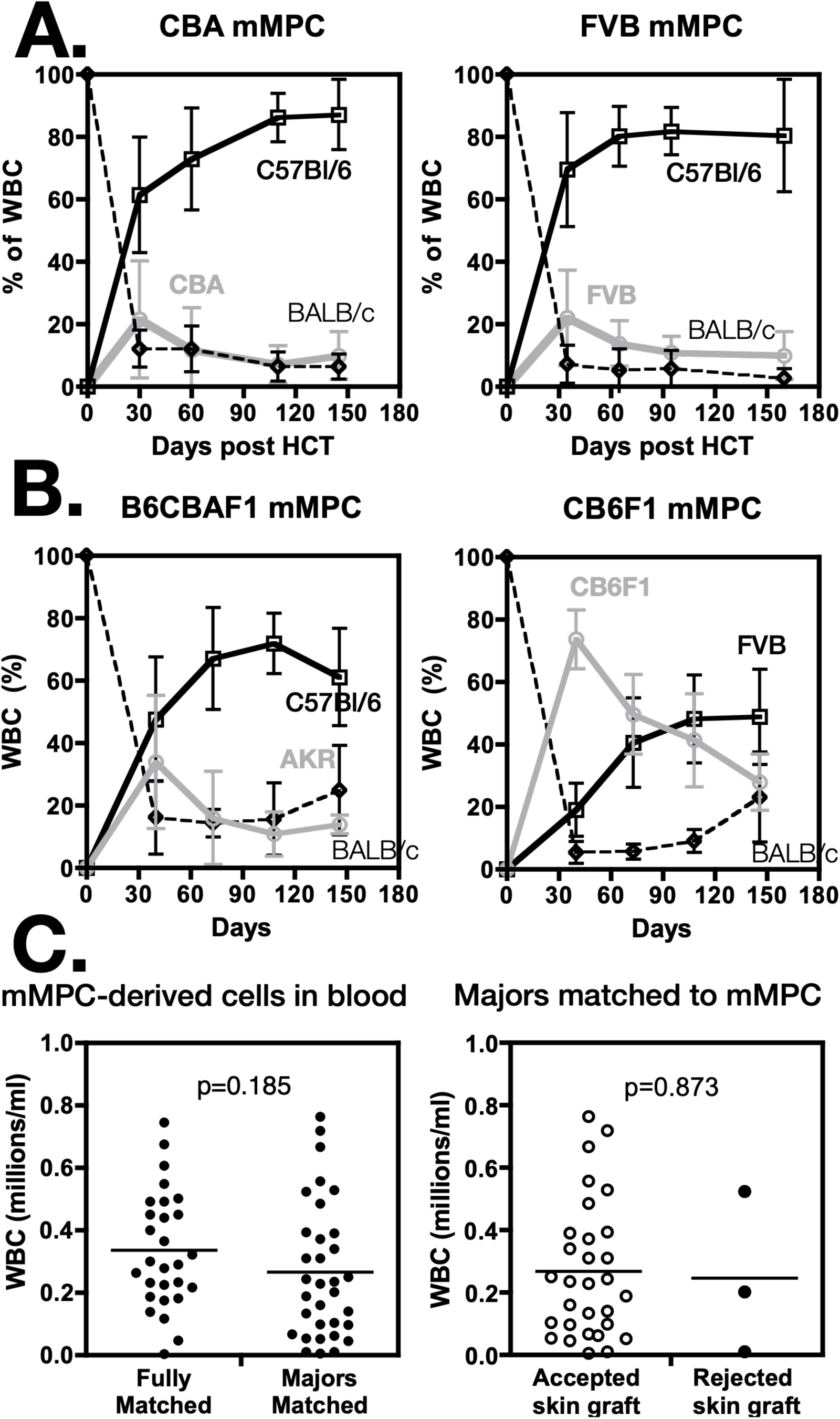
Blood analysis after reconstitution. (A) Blood composition of BALB/c host-derived (dashed line, diamonds), HSC-derived (solid black line, squares) and mMPC-derived cells (solid grey line, circles) at various times after HCT. HSC-derived cells are dominant. (B) Similar data for two experiments with F1 mMPC shows similar kinetics, albeit more robust engraftment, esp for CB6F1. Mean data and SD from 12 to 32 mice per time point, except for the 150-day time point of B6CBAF1 (9 mice). (C) The left plot shows the number of mMPC-derived cells in circulation at two months post hematopoietic/skin graft transplant for the listed graft types. The right plot compares the percentage of mMPC-derived cells in circulation two months after transplant in accepted versus rejected major-matched skin grafts.

The reconstitution data in Figure 4A-B are independent of skin graft matching and outcome. To test whether reconstitution was influenced by the skin grafts, we determined the absolute mMPC-derived cell numbers in circulation at 2 months post administration (Figure 4C). There was no significant difference (p=0.185, Fisher’s) between mice given fully matched skin grafts and mice given skin grafts only matched in the major MHC antigens of mMPC. As several skin grafts were rejected in the groups matched for major-antigens-only, we compared mMPC-derived cell numbers at two months following transplant between mice that accepted or rejected their skin grafts, pooling data from mice reconstituted with FVB and CBA mMPC. The average number of mMPC-derived cells does not differ significantly between the groups (p=0.873).

We also analyzed the presence of mMPC-derived cells in more detail for the entire period. Mice given CBA (Figure 5A) or FVB (Figure 5B) mMPCs did not clearly differ at any time in the percentage of mMPC-derived, HSC-derived, and host-derived cells in circulation between animals receiving mMPC that were fully matched, partially matched, or fully mismatched to the skin grafts (CBA mMPC-group only) at the time of hematopoietic transplant. Analysis of the main types of cells present in circulation showed no correlation between myeloid cells (Gr-1^+^ and/or CD11b^+^), B cells (CD45R^+^) and T cells (CD4^+^ or CD8^+^) at 30 days after hematopoietic and skin transplant, when the skin grafts start to get rejected, and the matching status (and rejection status) of skin grafts and mMPC grafts.

**Figure 5.**
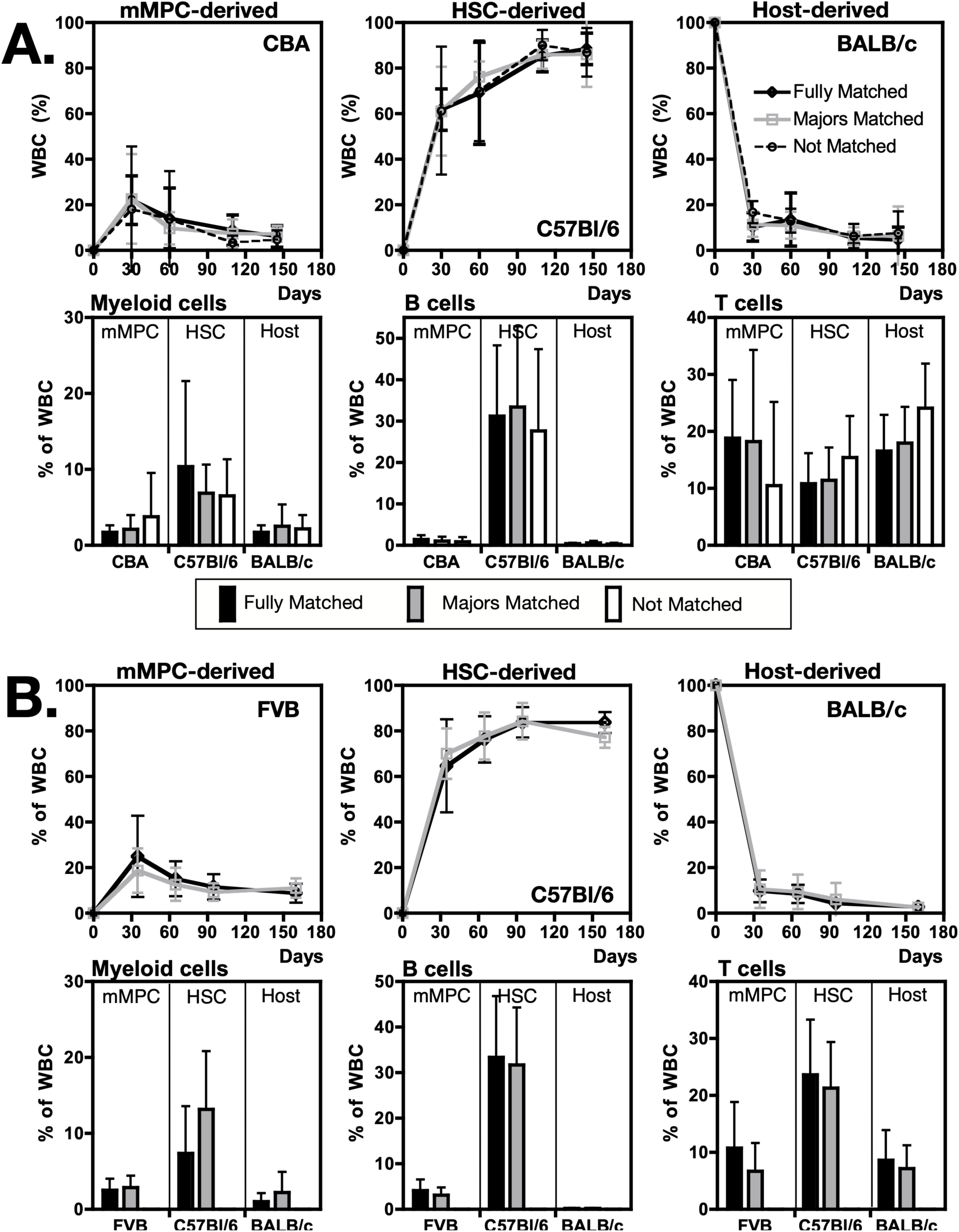
Blood analysis with different types of skin grafts. The figure shows mMPC-derived, HSC-derived, and host-derived cells in mice that received skin grafts fully matched to the mMPC, matched only for the major transplantation antigens, or not matched at all (panel A only). (A) BALB/c mice (H-2d) received 5×10^5^ CBA mMPCs (H-2k), 6×10^3^ C57Bl/6 HSCs (H-2b) and fully matching (CBA), majors only matched (AKR or C3H), or non-matched (FVB, H-2q) skin grafts. Top row: contribution of cells derived from each source in peripheral blood. Second row: percentage of myeloid cells (Gr-1^+^ and/or CD11b^+^), B cells (CD45R^+^), and T cells (CD4^+^ or CD8^+^) 30 days after transplantation. (B) BALB/c mice (H-2d) received 5×10^5^ FVB mMPCs (H-2q), 6×10^3^ C57Bl/6 HSCs (H-2b), and fully matching (FVB) or majors only matched (DBA) skin grafts. Top row: contribution of cells derived from each source in peripheral blood. Second row: percentage of myeloid cells (Gr-1^+^ and/or CD11b^+^), B cells (CD45R^+^) and T cells (CD4^+^ or CD8^+^) 30 days after transplantation. Mean data and SD from (A) 7 to 25 mice or (B) 8 to 16 mice per time point.

### Protection of heterotopic trachea grafts by mMPC

In addition to skin grafts we tested the ability of mMPC to protect trachea grafts placed under the skin (heterotopic) from rejection. BALB/c mice received FVB HSC and C57Bl/6 mMPC, after two months three trachea grafts were placed, one each originating from FVB (HSC-matched), C57Bl/6 (mMPC-matched), and CBA (mismatched) strains. The grafts were harvested and analyzed after one month. This is a transplant model for obliterative bronchiolitis following lung transplant that has been used extensively [20–22]. Using established criteria for this model such as swelling (reduced airway), lymphocyte infiltration and status of the airway epithelium, the results establish that unmatched trachea-grafts elicited a rejection response, while mMPC-donor matched tracheas were protected to a similar extent as HSC-donor matched tracheas (Figure 6).

**Figure 6.**
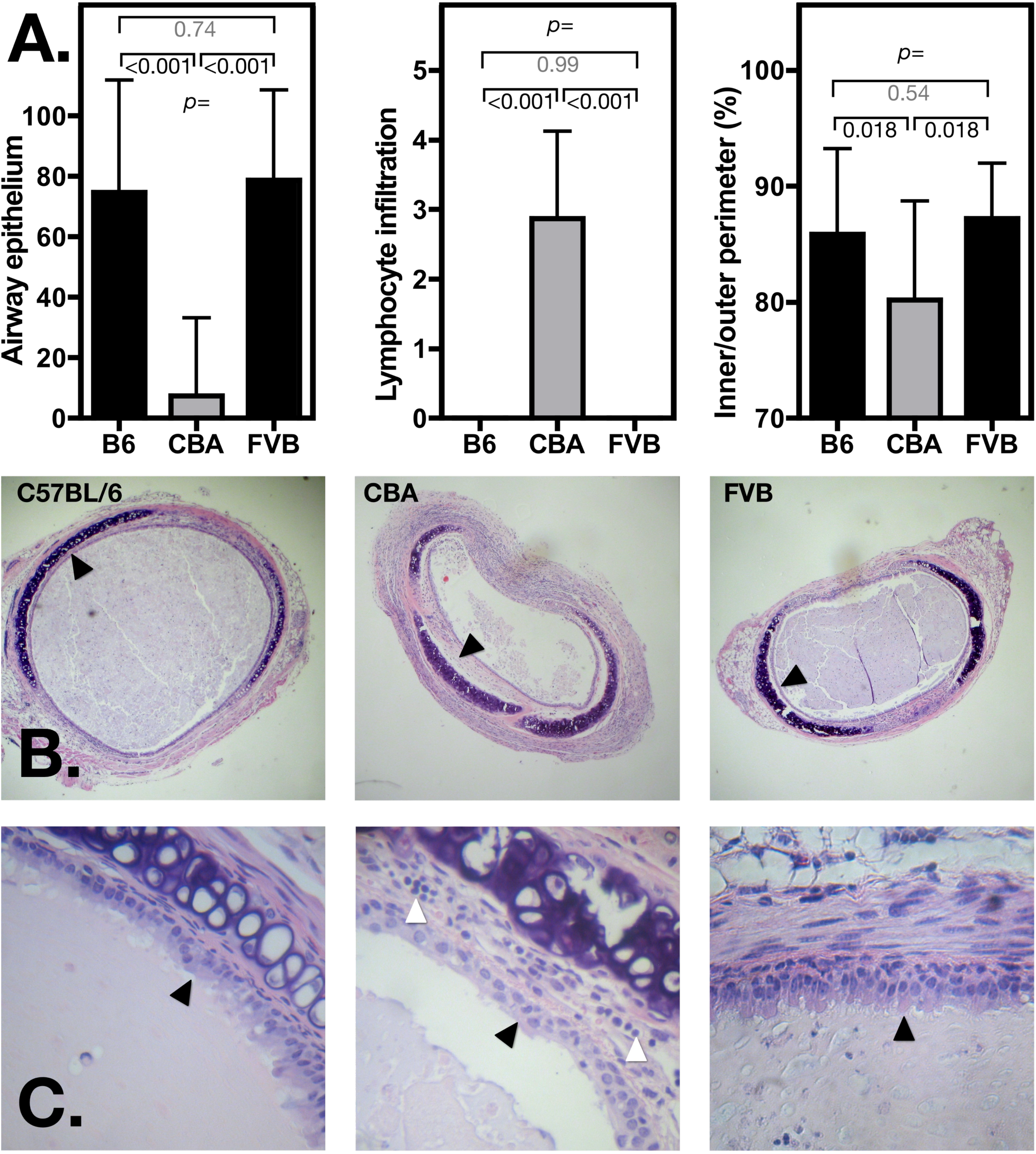
mMPC can protect matched heterotopic trachea transplants. mMPC-matched heterotopic trachea grafts (C57BL/6) are protected to a similar extent as HSC-matched (FVB) grafts. BALB/c mice were reconstituted with 4×10^3^ FVB HSCs and 5×10^5^ C57Bl/6 mMPCs. Trachea grafts (one each for C57BL/6, CBA, and FVB) were placed two months after reconstitution and analyzed one month later. (A) Airway epithelium growth (% of inner surface covered), lymphocyte infiltration (arbitrary units from 1-5), and ration of inner to outer perimeter were measured for each of the three grafts (n = 15). (B-C) Histopathology demonstrates that non-matched grafts (CBA) show major damage from rejection responses compared to C57BL/6 and FVB grafts, including (B) swelling (middle panel black arrow), loss of airway epithelium (middle panel black arrow), and (C) lymphocyte infiltration (middle panel white arrow).

### Non-lethal preconditioning

Lethal irradiation, while effective in preclinical models, is not the preferred preconditioning method clinically. We have tested a number of non-lethal preconditioning methods that result in robust immunodepression. We found that a combination of anti-thymocyte serum (ATS), anti-CD3, busulfan and rapamycin, administered over a two-week period as depicted in Fig 7A is not lethal and results in similar mMPC engraftment patterns as those seen following lethal irradiation (albeit that higher mMPC cell numbers need to be administered) and reliable protection of mMPC matched skin grafts (Figure 7). Importantly, we found that mMPC can be administered at least two weeks after skin graft placement. Overall, 31/34 mMPC matched grafts survived more than 5 months, which does not differ significantly from the protection seen for fully matched grafts in lethally irradiated mice (Fig 2), p=0.1 (logrank). We also established, using mitomycin-C treatment, that mMPC need to be mitotically active in order to be effective (Fig 7d).

**Figure 7.**
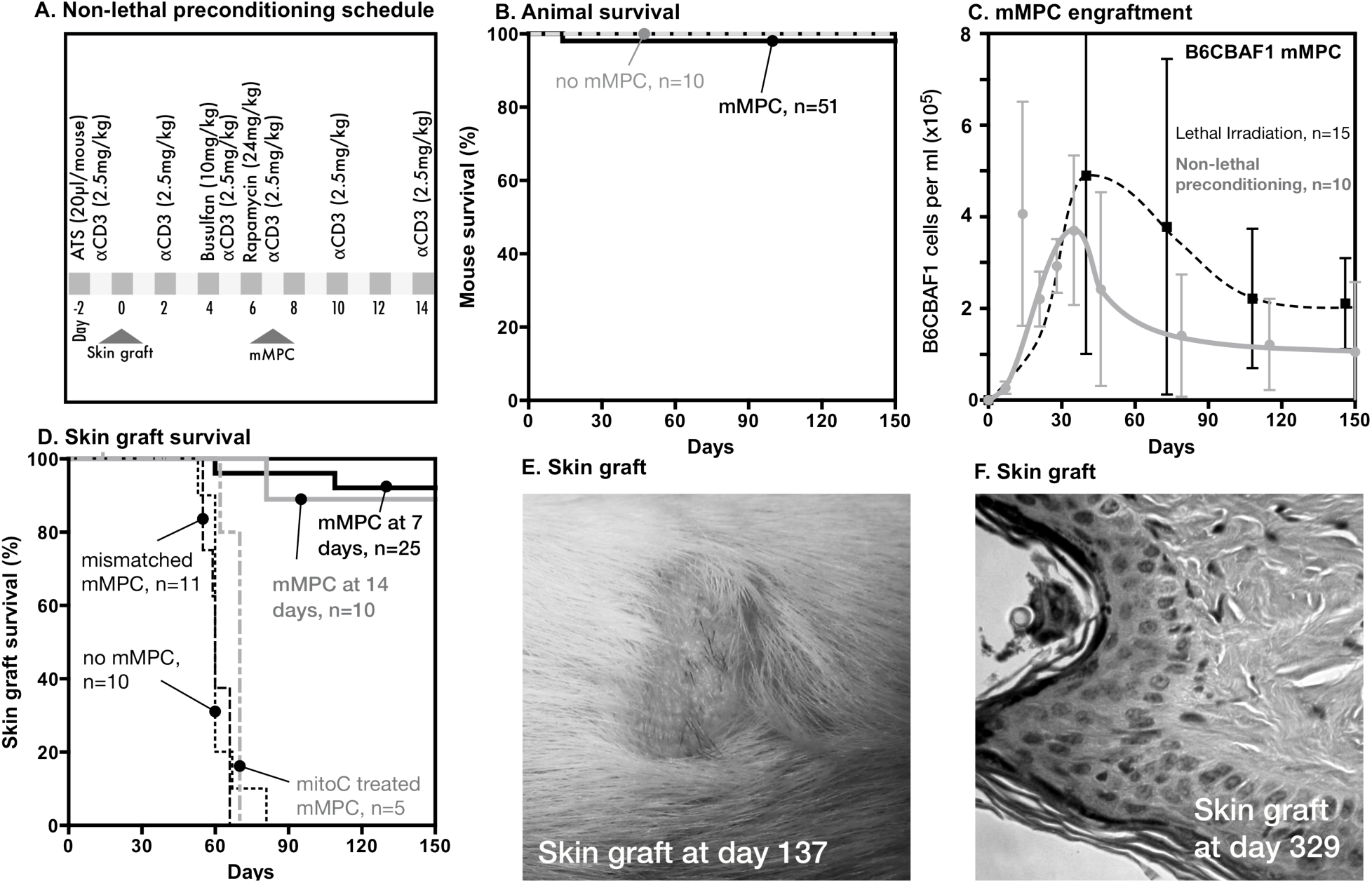
Non-lethal preconditioning. (A) The pre-conditioning protocol used, in combination with mMPC administration, to prevent rejection of mMPC matched skin grafts. (B) 5 month survival of 61 animals treated with this protocol, including 10 mice that received no mMPC or other hematopoietic graft. (C) mMPC engraftment levels in mice treated with lethal irradiation preconditioning and 5×10^5^ B6CBAF1 mMPC (dashed line, data from 2 experiments) and mice preconditioned using the protocol shown in panel A. These mice received 6×10^6^ B6CBAF1 mMPC (grey line, data from 2 experiments). Means and SD. (D) Skin graft survival in mice preconditioned as shown in panel A. All mice (other than the no-mMPC group) received between 3 and 6 million B6CBAF1 or CB6F1 mMPC. Black line shows survival of matched (C57Bl/6) skin grafts (data from 5 experiments). Grey line: similar, but the mMPC were given 14 days after skin graft placement (data from 2 experiments). Dashed black line mice receiving mismatched skin grafts (FVB) (data from 3 experiments). Dotted black line shows skin graft survival in mice receiving C57Bl/6 skin graft and preconditioning but no mMPC (data from 2 experiments). Dashed grey line shows graft rejection in mice receiving 3×10^6^ mitomycin C treated B6CBAF1 mMPC and C57Bl/6 skin grafts (data from 1 experiment). (E) Gross morphology and histology of an mMPC-matched skin graft placed using non-lethal preconditioning.

### T and B cell responses

To start determining the mechanisms that allow mMPC to prevent rejection of matched grafts, we investigated T cell responses using MLR, and investigated the presence of H-2 specific antibodies using the cells and serum from regular bleeds of mice from three independent experiments. As depicted in Fig 8A-B, we found significant increases in anti-skin graft antibodies for FVB skin grafts not matched to mMPC during rejection (confirming a functioning B cell response), but no increase in antibodies recognizing C57Bl/6 skin grafts in mice that received C57Bl/6 mMPC. Fig 8C shows that both FVB and C57Bl/6 skin grafts elicit an antibody response in mice that did not receive mMPC. Fig 8D, E shows that T cells from these animals have a significantly reduced response to mMPC matched stimulator cells compared to non-matched cells. This clearly suggests that at least some T cell response has returned following the preconditioning.

**Figure 8.**
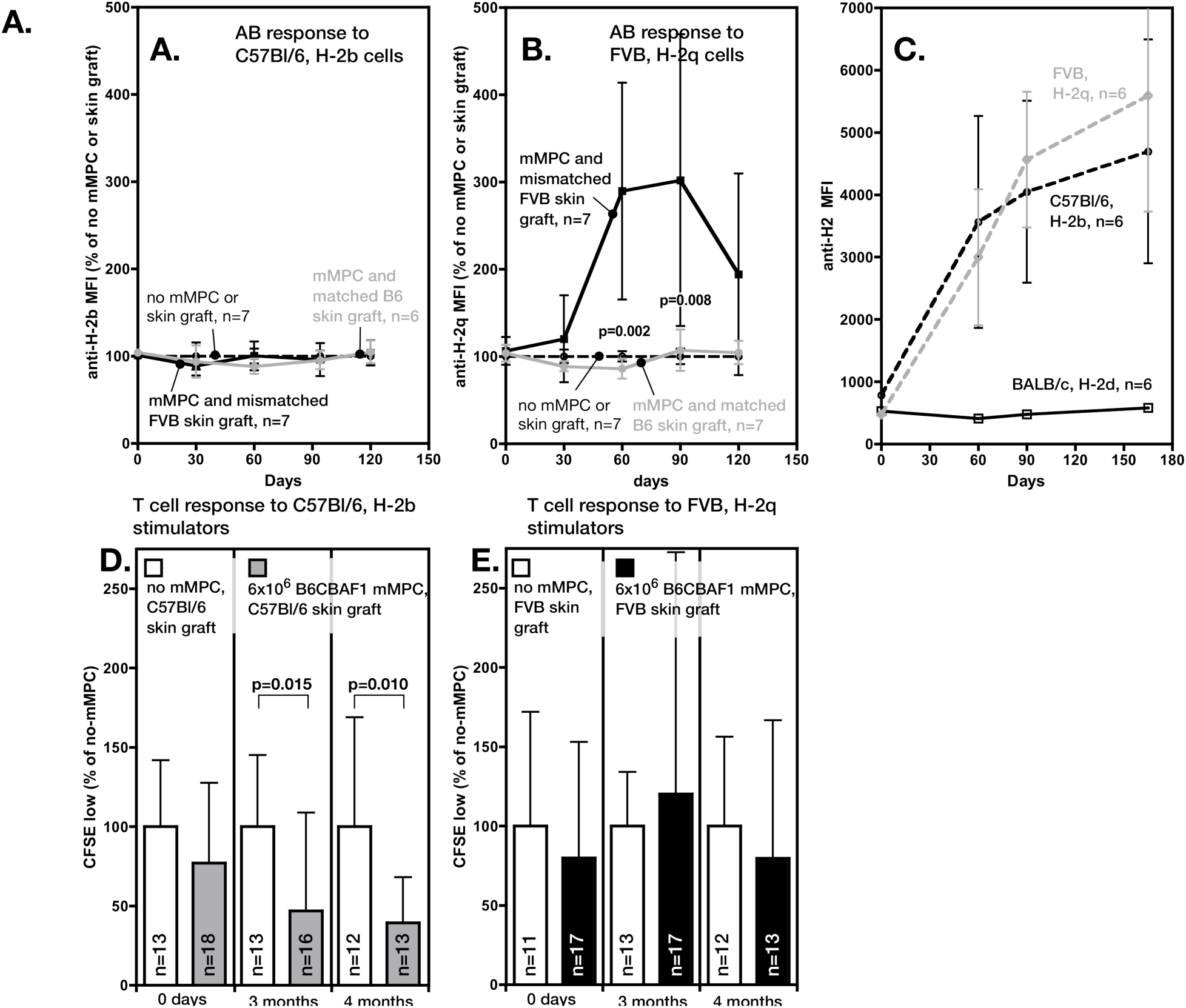
B and T cell responses. (A) Antibody levels in plasma at different times after mMPC administration as shown in Fig 7a. Data relative to that of mice that received non-lethal preconditioning, but neither mMPC, nor skin grafts. BALB/c mice received 6×10^6^ B6CBAF1 mMPC and either matched C57Bl/6 or mismatched FVB skin grafts. Groups as indicated. There is no difference in circulating antibodies recognizing C57Bl/6 cells in animals that have received mMPC. (B) Levels of circulating anti-H-2q antibodies in mice that received mMPC-mismatched FVB skin grafts significantly increases during rejection of these grafts (approx 3 months after placement) (C) Mice that received non-lethal preconditioning but no mMPC reject both C57Bl/6 and FVB skin grafts, and show similar increases in anti-H-2b and anti-H-2q antibodies in circulation. No increase in antibodies recognizing host tissues (H-2d) is seen. (D) shows that the T cell response against C57Bl/6 stimulator cells is reduced in mice receiving B6CBAF1 mMPC at 3 and 4 months post transplant. There is no similar reduction in the response to FVB stimulation (E). In both the response is shown relative to that of mice not receiving cells. Data from 3 experiments. Means and SD, p-values are shown where less than 0.05.

## Discussion

The long-term survival of transplanted organs remains limited [23] and new treatment options are needed [1, 2, 9]. High-level HCT has held up in both experimental and clinical settings, but has been limited by the associated complications. It continues to be actively pursued for tolerance induction in solid organ transplant recipients [24–26]. We have demonstrated that allogeneic myeloid progenitor cells efficiently prevent rejection of MHC-matched tissue grafts that have been placed upto two months after preconditioning/MPC administration, after hematopoietic cell levels have normalized to a large extend [13, 19]. Specifically, we have reported that adding third party MPCs to either an allogeneic or autologous HSC transplantation results in robust protection for MPC-matched skin grafts, without long-term high-level engraftment of MPC-derived cells. Importantly, a clinical grade human MPC product (romyelocel-L) has been developed to prevent bacterial and fungal infections in patients with prolonged neutropenia and has shown an acceptable safety profile in 151 subjects over three clinical trials, facilitating translation of graft protection to the clinic.

Myeloid lineages contain regulatory cells such as myeloid-derived suppressor cells [27, 28], tolerogenic DC [29, 30] and M2-polarized macrophages, including tumor-associated macrophages [31–33]. All have been implicated in tolerance induction for solid organ transplantation and several are being tested in either phase I clinical trials or at the preclinical stage [8, 18, 34, 35]. Culture derived MPCs are able to differentiate into all of these populations which may play overlapping or partially redundant roles in preventing skin graft rejection.

There are practical limitations in devising parameters for the clinical use of myeloid progenitor cells. One variable tested here is the degree of matching required between the MPC and the skin graft. These experiments clearly established that while minor mismatches do not significantly reduce the number of rejected grafts, haploidentical skin grafts, with one mismatched allele to mMPC, HSC, and host are not protected. Grafts are only protected if each MHC allele is matched to an MHC allele in the mMPC, host or HSC donor. Because minor transplantation antigen mismatches are tolerated it should be possible to produce mMPCs based on matching major transplantation antigens (MHC) rather than from the fully matched organ donor itself. The fact that MHC molecules present on different cells (mMPC and host) efficiently protects grafts as depicted in figure 2C suggests that pooled donor batches of mMPC will be effective in providing coverage for an extended set of haplotypes.

We did not find evidence that either the degree of matching between mMPC and skin graft or the rejection status correlated with the numbers of mMPC-derived cells in circulation, confirming that cell numbers in circulation are not a good predictor of graft protection in this model. It will be interesting to determine if mMPC-derived cell numbers in the graft, or in other tissues such as thymus, bone marrow or secondary lymphoid organs, are a better indicator.

We have also established that mMPC can protect skin grafts when using a non-lethal preconditioning protocol, which abrogates the need for a simultaneous HSC transplant and removes a major barrier to potential clinical use. However, higher numbers of mitotically active mMPC are needed. This protocol allows for mMPC administration at least 2 weeks post skin graft placement, enough time to expand organ donor bone marrow cells into MPC and use fully matched MPC cytotherapy.

Our results demonstrate that mMPC MHC needs to be closely matched to the organ donor MHC. Pooling MPC from 10 donors would increase the number of potential MHC alleles in a batch to 20 for each gene, given the absence of allelic exclusion. As an alternative, fully matched donor-specific MPC expanded from individual living or deceased organ donors, are possible using the non-lethal preconditioning protocol. The rejection of third party (unmatched) skin and trachea grafts, as well as the antibody response in preconditioned mice to non-MPC matched targets (this paper), as well as the ability of mMPC to protect transplantation antigen-matched skin grafts when placed two months after preconditioning/mMPC transplant (REF) indicate skin graft protection while immune competence improves, arguing for mMPC-induced tolerance for matched grafts. Further optimization of the preconditioning protocol combined with further characterization of the return to immune competence in these animals will be necessary and important parameters for developing clinical use scenarios.

## Acknowledgements

Supported by grants from NIAID, 1R41AI108016-01, and a Katherine B Richardson Endowment Award. AS and TF were employed by Cellerant Therapeutics at the time this research was conducted. The authors otherwise have no conflicts to disclose.

## Abbreviations

DC: Dendritic Cell
HCT: Hematopoietic Cell Transplantation
HSC: Hematopoietic Stem Cell
MHC: Major Histocompatibility Complex
MPC: Myeloid Progenitor Cell
mMPC: Mouse Myeloid Progenitor Cell derived by ex vivo expansion

